# Ballistic food approaches in *Parhyale hawaiensis* require the antennae

**DOI:** 10.64898/2026.06.16.732625

**Authors:** Theresa Steele, Katherine Nagel

## Abstract

Many arthropods (insects and crustaceans) rely on their antennular chemosensory system to detect key environmental resources like food. While odor mediated food search is well studied in insects, characterization of crustacean chemosensory behavior has been limited by the long lifespans and large size of traditional crustacean model species. Here, we report the first characterizations of the food search behaviors of the genetically tractable amphipod crustacean, *Parhyale hawaiensis*. We find that *Parhyale* can locate an odorous food pellet, and predominantly approach food using direct, targeted swims from the arena walls. Removal of both first and second antennae dramatically reduced foraging success and impaired Parhyale’s ability to control take-off angle and maintain a stable heading during swims. Removal of the first or second antenna alone did not significantly disrupt foraging, and resulted in mild disruption of orientation phenotypes. Intact animals performed sharp turns near the location of the food pellet, which were observed when either first or second antenna were present, but not when all antennae were removed. Turns were longer and had higher average angular velocities following removal of either set of antennae, with full antenna removals representing the most extreme phenotype. In contrast with the long-held theory that the crustacean second antennae exclusively mediate contact chemosensation, we report that first- and second- antennae both contribute similarly to food localization and stabilization of locomotion in Parhyale in our behavioral paradigm. This work establishes Parhyale as an accessible model for studying olfactory behaviors in an aquatic arthropod.

## Introduction

The olfactory navigation behaviors of arthropods are a powerful system for studying behavioral adaptation across environmental niches. Many arthropod species use odor as a long-range cue of salient environmental features such as food and mates, including fruit flies (Stupski and van Breugel 2024; Álvarez-Salvado et al. 2018), moths (Kennedy et al. 1981; Willis et al. 2013), blue crabs (Page et al. 2011a, 2011b), lobster (Basil and Atema 1994; Horner et al. 2004), and crayfish (Kraus-Epley and Moore 2002). However, olfactory navigation strategies must adapt to both an animal’s biology (such as the structure of olfactory sensory appendages and an animal’s locomotor modalities), as well as the structure of relevant odors in the environment (e.g. the constancy or intermittency of odor signals). For example, while a walking carpenter ant may use its articulating antenna to probe the environment and actively compare odor over space to follow the static edges of an odor trail (Draft et al. 2018), a crawling *Drosophila* larva must stop to perform active head-sweeps in order to sample olfactory gradients with its cephalic dorsal organ, integrating odor encounters over time to drive navigation (Gomez-Marin et al. 2011; Gomez-Marin and Louis 2014). Studying the olfactory navigation strategies employed by arthropods across olfactory environments may provide an accessible model for understanding behavioral diversification. In particular, comparing the navigation strategies of aquatic and terrestrial arthropods is of particular interest due to fundamental differences in what constitutes an odorant, and how odors are dispersed across these environments (reviewed Steele et al. 2023; Krieger et al. 2021).

Crustaceans exhibit robust olfactory behaviors driven by a complex chemosensory system. Olfactory navigation behaviors have been observed in many crustaceans, including blue crabs (Page et al. 2011a, 2011b), lobster (Basil and Atema 1994), and crayfish (Wolf et al. 2004; Kraus-Epley and Moore 2002). Like insects, large decapods have been shown to use both flow and odor cues to locate odorous food (Weissburg and Dusenbery 2002; Reeder and Ache 1980). Unlike insects, crustaceans possess two sets of antennae: the smaller, dorsally located first antennae (or ‘antennules’), and the larger, ventral second antennae. While both antennae possess bimodal chemo- and mechanosensory sensilla, only the first antenna possess sensilla containing olfactory sensory neurons that project to glomeruli in the antennal lobe (Kümmerlen et al., 2025) leading a proposed distinction between “olfaction” mediated by unimodal sensilla and “distributed chemoreception” mediated by bimodal sensilla (reviewed Harzsch and Krieger 2018; Krieger et al. 2021; Loesel et al. 2013). Selective ablation experiments have demonstrated that bimodal sensilla of the first antenna are sufficient to mediate navigation behaviors in decapod crustaceans (Horner et al. 2004). However, a behavioral role for the bimodal sensilla of the second antennae has not been clearly identified. Detailed characterization of the contributions of each set of antennae to olfactory navigation strategies in crustaceans has historically been limited to low sample sizes due to the large size of traditional decapod models.

The amphipod crustacean *Parhyale hawaiensis* (Dana, 1853) is an emerging genetic model well suited to comparative studies (Louis et al. 2026; Kwiatkowski et al. 2023). *Parhyale* are a small (∼1 cm in body length) and hardy model system with established techniques for husbandry (Paris et al. 2022), novel transgenesis in targeted cell lines, (Pavlopoulos and Averof 2005; Forbes et al. 2026), and targeted mutagenesis (Martin et al. 2016). Neuroanatomical work has extensively characterized the brain anatomy and olfactory systems of *Parhyale*, identifying approximately 13,000 neurons both male and female brains (Wittfoth et al. 2019) and approximately 50 olfactory glomeruli innervated by ∼1200 olfactory sensory neurons (Kümmerlen et al., 2025). Drosophila have served as a highly effective model for studying olfactory behaviors due to the simplicity of their olfactory systems, and the development of transgenic tools to allow investigation into the function of populations of neurons within olfactory and navigation circuits. This easy cultivation, genetic accessibility, and relatively simplified olfactory system all position *Parhyale* well as a model for studying olfactory behaviors and circuit function in crustaceans.

In this study, we characterize the food search behaviors of freely swimming *Parhyale*. We constructed an imaging apparatus and developed a novel behavioral analysis pipeline to quantify locomotion in the presence of a food pellet and a wax sham pellet. The majority of *Parhyale* were able to locate food, which was largely abolished by ablation of both the antennae and antennules. Intact *Parhyale* made ballistic, directed approaches to the food pellet, often following repeated swims through the arena. Antennae removal impaired both the ability to make aimed take-offs from the wall, and to maintain a consistent heading during swimming.

Removal of single antenna pairs demonstrated that both the first and second antenna alone were sufficient for successful food search behaviors, despite causing moderate impairment to locomotion. First- or second antennae removal resulted in a small increase in heading instability during swimming. Features of behavior in first or second antenna removed animals were consistently similar, and intermediate between the behavior of intact and full antenna removed animals. We additionally examined turning behaviors and found that intact animals exhibited sharp turns biased to an approximately 3 cm radius around the food pellet. While full antenna removal disrupted this pattern, first or second antenna removal did not. All antenna removals altered turn structure, causing turns to have longer durations and higher angular velocities. Overall, these findings suggest that (1) *Parhyale* use a ballistic food search strategy that is distinct from gradient ascent strategies found in other arthropods (2) antennae are essential for successful food search and stable locomotion and (3) that first and second antennae can redundantly facilitate food localization in low flow conditions.

## Materials and methods

### Animals

Behavioral experiments were performed with an inbred strain of *Parhyale hawaiensis*, Chicago F (Paris et al. 2022; Parchem et al. 2010). *Parhyale* colonies were maintained at room temperature under natural light in 35% seawater in 5.2-gallon bins (McMaster 6686T73), with a coral and sand substrate (Bio-Activ Live Aragonite, CaribSea Arag-Alive Sand). Colonies were fed weekly with sinking fish food (Aqueon,100552011) and dried kelp flakes.

We imaged behavior in both intact *Parhyale* (N=49), and in *Parhyale* with one or both sets of antennae removed (N=37 first antennae removed, N=37 second antennae removed, N =39 both antennae removed). To remove antennae, animals were anaesthetized by being placed directly into ice cold sea water, and antenna were removed at the end of their initial segment (supplemental figure 1) using fine sharpened forceps.

Both intact and antenna-removed *Parhyale* were food-deprived for 5-17 days prior to behavioral experiments (average starvation period intact: 9-17 days, mean = 12.3; full antenna removal: 5-19 days, mean =12.4 days; second antenna removal (2^nd^ antx) 9-17 days, mean = 13.7 days; first antenna removal (1^st^ antx), 8-17 days, mean = 12.4 days. Experimental animals were isolated from the colony and starved in groups of 10-20 in small plastic containers (McMaster 4494T22) which were bubbled continuously with oxygen. Plastic garden mesh was provided as a substrate for animals to latch onto. For intact animals, 40/49 were placed directly from the rearing aquarium into food depravation chambers, while 9/49 were anaesthetized and food deprived alongside antenna-removed *Parhyale*. Antenna removals were verified by visual inspection of *Parhyale* prior to experimentation to exclude animals exhibiting antennal regrowth.

### Experimental Apparatus

We designed a custom behavioral imaging arena to monitor *Parhyale* behavior (Supp Fig. 1). The arena consisted of layers of laser cut acrylic of varying thicknesses (see below) to form a reservoir, and laser cut rubber seals to maintain water tightness, which were designed in Adobe illustrator (design: Adobe Systems, San Jose, CA; plastics: Pololu Corp, Las Vegas, NV and McMaster, Robbinsville, NJ; laser cutting: Pololu). The internal layers of the arena consisted of 1” thick acrylic (McMaster 8560K322), .125” silicone gasket (McMaster 8525T184), and a bottom layer of .5” acrylic (McMaster 8560K266), which were held together using set screws (McMaster 92185A307).

**Figure 1.**
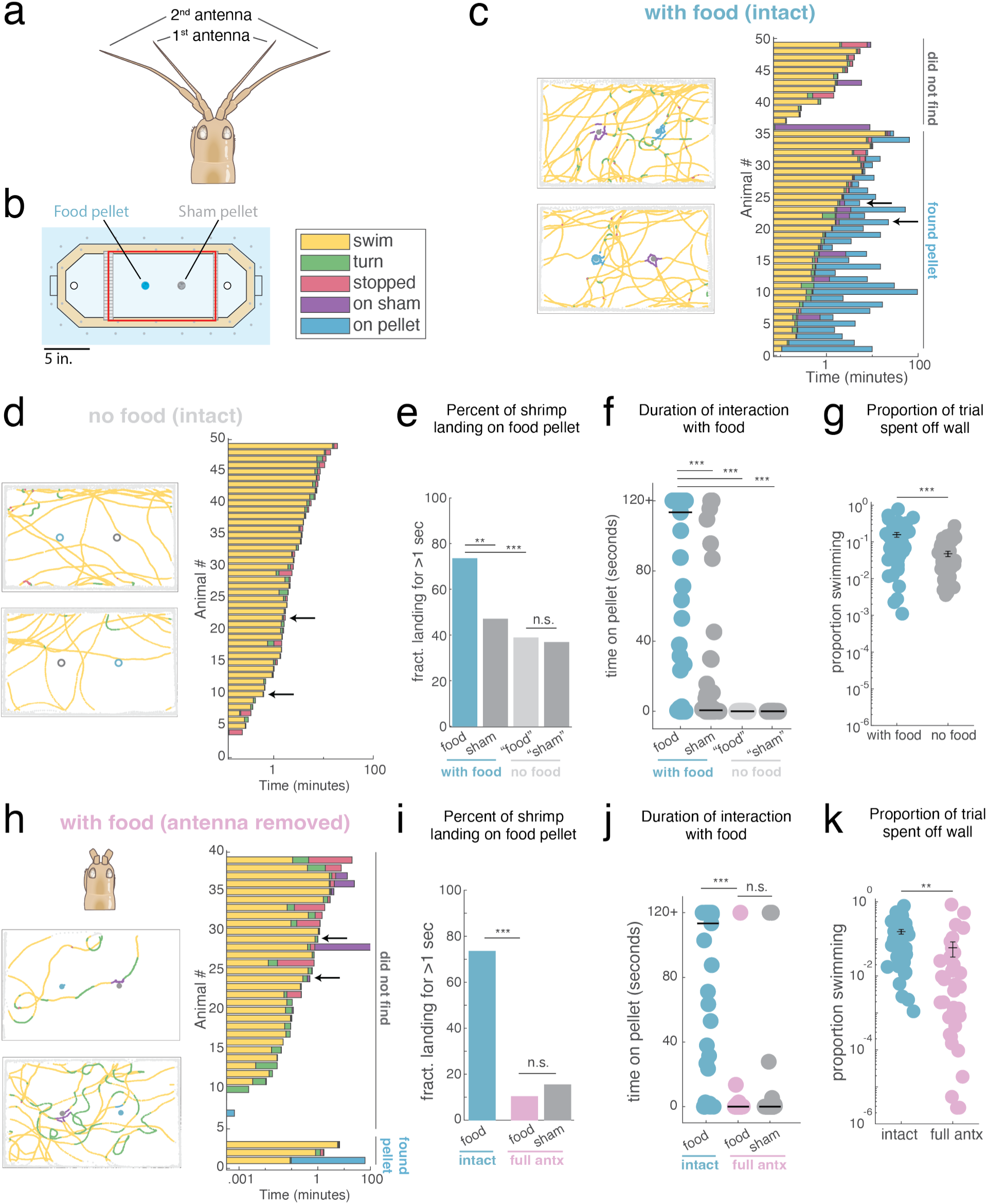
*Parhyale* exhibit behavioral responses to food that are mediated by the antennae. a. Illustration of the antennal system of *Parhyale hawaiensis*. b. (left) Schematic of behavioral recording apparatus. (right) color code used for ethogram c. (left) examples of locomotion in intact animals in the presence of food, color coded by behavioral state (swim, turn, stop, sham, pellet, wall walking); (right) ethogram of active behaviors (excluding wall walking) in the presence of food for all intact experiments (n= 49), arrows indicating experiments used as examples. d. (left) examples of locomotion in intact animals during no-food control trials, color coded by behavioral state (swim, turn, stop, sham, pellet, wall walking); (right) ethogram of active behaviors (excluding wall walking) during no-food control trials, for all intact experiments (N = 49), arrows indicating experiments used as examples. Symbols show locations of pellet and sham on associated food trials. e. Percent of animals landing on the food and sham pellet for intact with and without food conditions. For no food controls, “on-pellet” or “on-sham” times were determined using the position of the food and sham pellets in the corresponding experimental trial (ChiSquare, with food pellet vs sham p = .0073, with food pellet vs no food “pellet” p = .00054, no food “pellet” vs no food “sham” p = .834) f. Duration of longest interaction with the food or sham pellet for intact animals with and without food. Black line indicates median duration: 113.4 seconds for food compared to 0.6 sec for sham (rank sum p= 1.76 e-5), and median duration of 0 seconds for control “pellet” or “sham” (p = 1.196 e-12 relative to food) g. Proportion of trial spent in active behaviors (those other than wall walking). Intact with vs without food ANOVA p = 0.00016) h. (left) examples of locomotion in animals with all antennae removed in the presence of food, color coded by behavioral state (swim, turn, stop, sham, pellet, wall walking); (right) ethogram of active behaviors (excluding wall walking) in the presence of food for all full antenna removed animals (N = 39), arrows indicating experiments used as examples. i. Percent of animals landing on the food pellet for intact and full antenna removed conditions, and on the sham pellet for the full antenna removed condition. (ChiSquare, intact pellet vs antx pellet p = 3.3 e-9; antx pellet vs antx sham p = .4982) j. Duration of longest interaction with the food or sham pellet for intact animals with and without food. Black line indicates median duration: 113.4 seconds for intact compared to 0 sec for antenna removed approach to food (rank sum p= 1.59 e-9), and median duration of 0 seconds for antenna removed approaches to food or sham (p =.3595) k. Proportion of trial spent in active behaviors (those other than wall walking, intact vs antx with food rank sum p = .0019).

Water was provided to the experimental arena using a recirculating system with an ozone filter (supplemental fig 1a). 35% sea water (Instant Ocean) was stored in a sump tank (Trigger Systems Crystal Sump 30, SKU : 209219) and continuously filtered through an ozone and charcoal filtration system (Avast Aquatics Mutiny Ozone Reactor V2.0, Avast Aquatics Mutiny blend granular activated carbon, Ozotech Poseidon 200 ozone generator) to remove organic materials, while oxidation reduction potential was monitored with a ORP controller (Milwaukee Instruments MC510). Sea water was pumped from the sump tank to the behavioral arena using a DC controllable return pump (Reef Octopus VarioS-2), and water level was maintained by using a custom 3D-printed standpipe fitted into a drain in the bottom of the arena.

The arena was mounted in an imaging chamber constructed out of 80/20 posts (McMaster: 47065T101) held in place with brackets (McMaster: 47065T236), on a breadboard base (Thorlabs). The rig was illuminated by infrared LED strips (850 nm) mounted on an aluminum sheet (McMaster: 88835K15) beneath the arena, with a sheet of diffusive plastic placed 5 cm above the light panel to provide even illumination (Eplastics: ACRY31430.125PM). *Parhyale* were imaged from above with a monochrome USB 3.0 camera (Basler: acA1920-155um) and a 12 mm 2/3’’ lens (Computar: M1214-MP2). LEDs were strobed at 50Hz to synchronize illumination with each camera frame using buckblock drivers (Digikey) and an Arduino microprocessor (teensy 2.0, PJRC). All experiments were performed in the dark, using infrared (IR) lighting and cameras to monitor behavior.

Imaging was performed with custom software written in Labview (National Instruments, Austin, TX), adapted from previously published work (Álvarez-Salvado et al. 2018). *Parhyale* position within the arena was calculated by subtracting a previously acquired background image, thresholding, and filtering to remove noise particles smaller than 25 pixels and larger than 150 pixels. The center of mass of the largest remaining particle was then calculated in real time and saved to disk, while a frame of 200×200 pixels around the centroid was saved as a video. Behavior and video were recorded in 2-minute trials, with approximately 3 seconds between trials.

Following image acquisition, we labeled saved image frames using DeepLabCut (Mathis et al. 2018) to extract the animal’s heading. We manually annotated 37 points on the shrimp: the tip, first joint, and base of each antenna and four prominent legs, both eyes, the 11 exoskeletal segments along the back of the animal, including the endpoint of the body (supplemental fig. 1b). We randomly selected 20 frames each from 32 videos for manual annotation and trained a ResNet-50 based model in DeepLabCut (version 2.0.5.1) using a training/test fraction of 0.8, meaning that of the annotated frames used to train the model, 20% were held in reserve to test model function. Following training, additional outlier frames were selected from to refine labelling output, up to a total of 862 frames. The network used achieved a training error of 6.42 and a test error of 10.38 using a p-cutoff of 0.7. Batch analysis of video was performed using custom Python 3.10.12 scripts.

### Analysis of behavioral data

All analyses were performed in Matlab (Mathworks, Natick, MA). Large jumps in centroid x or y position (>10 pixels in a single sample), were classified as tracking errors, and removed. x and y coordinates were converted to mm, and velocity was calculated as the displacement between subsequent coordinates multiplied by the frame rate (50 Hz). The location of the food and sham pellet were measured on the corresponding background image for each trial using the *ginput* function, and distance from the food / sham pellet were calculated as the hypotenuse of displacement between the animal and the food or sham pellet.

DeepLabCut annotations with a likelihood of p = .7 or higher were imported and integrated with centroid tracking data. Large jumps in annotation x or y position (> 10 pixels per frame) were removed from tracking, and annotation sets with <10 well tracked points were discarded for low quality. Small gaps in annotation (<25 frames) were filled with 1-dimensional interpolation using the interp1 function, and annotations were lowpass filtered at 2.5 Hz with a Butterworth filter. Heading was then calculated as the axis between the animal’s head point (an average of antenna base and eye positions) and tail point. The heading angle was unwrapped (phase corrected to be continuous, rather than circular) and angular velocity was calculated as the derivative of heading angle multiplied by the frame rate (50 Hz).

Data from individual recordings were concatenated using file timestamps to create continuous, real-time vectors of all measured parameters. From these, a unitless metric of instantaneous path curvature was calculated as the arc-chord ratio of paths over .5 seconds (25 samples).

#### Ethogram

To classify behavior at each time point, we created an ethogram based on features of locomotion. Periods in which the animal was within 1 cm of the wall position were labeled “wall walking”, and periods in which animals were within 1.25 cm of the food or sham pellet location were labeled as “on food” or “on sham” respectively. Because the walls, food pellet, and sham pellet could create tracking errors (by obscuring the position of the shrimp), and animals tended to engage in each behavior for extended periods, untracked segments between successive periods of wall walking, on pellet, or on sham behavior were categorized as the corresponding behavior. Segments where the instantaneous curvature was high (>1.1) and forward velocity was above a low baseline (>.5 cm/sec) were categorized as “turns”, and segments where forward velocity was low (<.5 cm/sec) were categorized as “stops”. All other behavior was considered “swimming”.

#### Swim analyses

Normalized swim lengths were calculated as the total path length during swimming/turning normalized to the total recording length (to account for differences in recording lengths). The proportion of the trial that animals were active was calculated as the duration of the trial in which the animal was tracked doing a behavior other than wall walking divided by the total duration of tracked behavior.

To identify swim bouts, sequences of behavior in between periods of wall walking, on pellet, on sham, and untracked behaviors were extracted, allowing for gaps up to 1 second to account for brief passes by the food/sham pellet or small tracking errors. For these swims, path deviation (the ratio of ground distance covered to direct start-end path length), average angular velocity, and the standard deviation of heading angle (based on unwrapped heading) and angular velocity were calculated as metrics of path instability.

#### Takeoff analyses

Takeoff events were classified as the moment in which an animal transitioned from “wall walking” to swimming/turning behaviors. Only takeoffs preceding swims that lasted >.5 seconds were considered in this analysis. Takeoff angles were calculated by rotating each trajectory by a rotation matrix depending on the wall of origin, and then takeoff angle was calculated based on the vector direction of the first .2 seconds following wall takeoff, ensuring that all takeoff angles were oriented from 0 to 180 degrees.

#### Food/sham/control approach analyses

Animals that exhibited “on food” behaviors that lasted longer than 1 second were classified as having found the food pellet. For these animals, food approaches were defined as the trajectory between the last departure from the wall before food was found, and the point at which animals reached 1.5 cm from the food pellet centroid. This excluded reorienting turns that often occurred at the end of food approaches, to allow for better comparison of the paths leading an animal to food. For display purposes, approaches were rotated to be aligned to the direction of travel at the final .2 seconds of approach. The same analyses were repeated for approaches to the sham pellet. For non-experimental trials, approaches to the locations in the arena where the food pellet was placed in the corresponding experimental trial were used to extract control “approach” behaviors.

#### Turn analyses

For analyses of turn location and locomotor features of turns, individual turns were identified based on ethogram labelling, allowing for small gaps (<.3 seconds) due to tracking errors or speed variability during turning. Turn location was calculated as the center of mass of each turn and turn distance from the sham/pellet was found using the average sham/pellet distance for the turn period. Angular velocity and path length during turns were calculates as previously described.

### Statistical Analyses

#### Jackknife Resampling

For the distributions in figures 2-6, error bars were calculated through Jackknife cross-validation. Parameter distributions were recalculated systematically leaving out measurements from each individual animal to create an n-by-m matrix (where n is the number of animals and m is the number of bins in the distribution). To calculate the bias of the distribution, we calculated the Jackknife standard deviation of the resampled distributions:

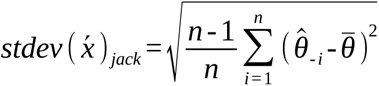

**Figure 2.**
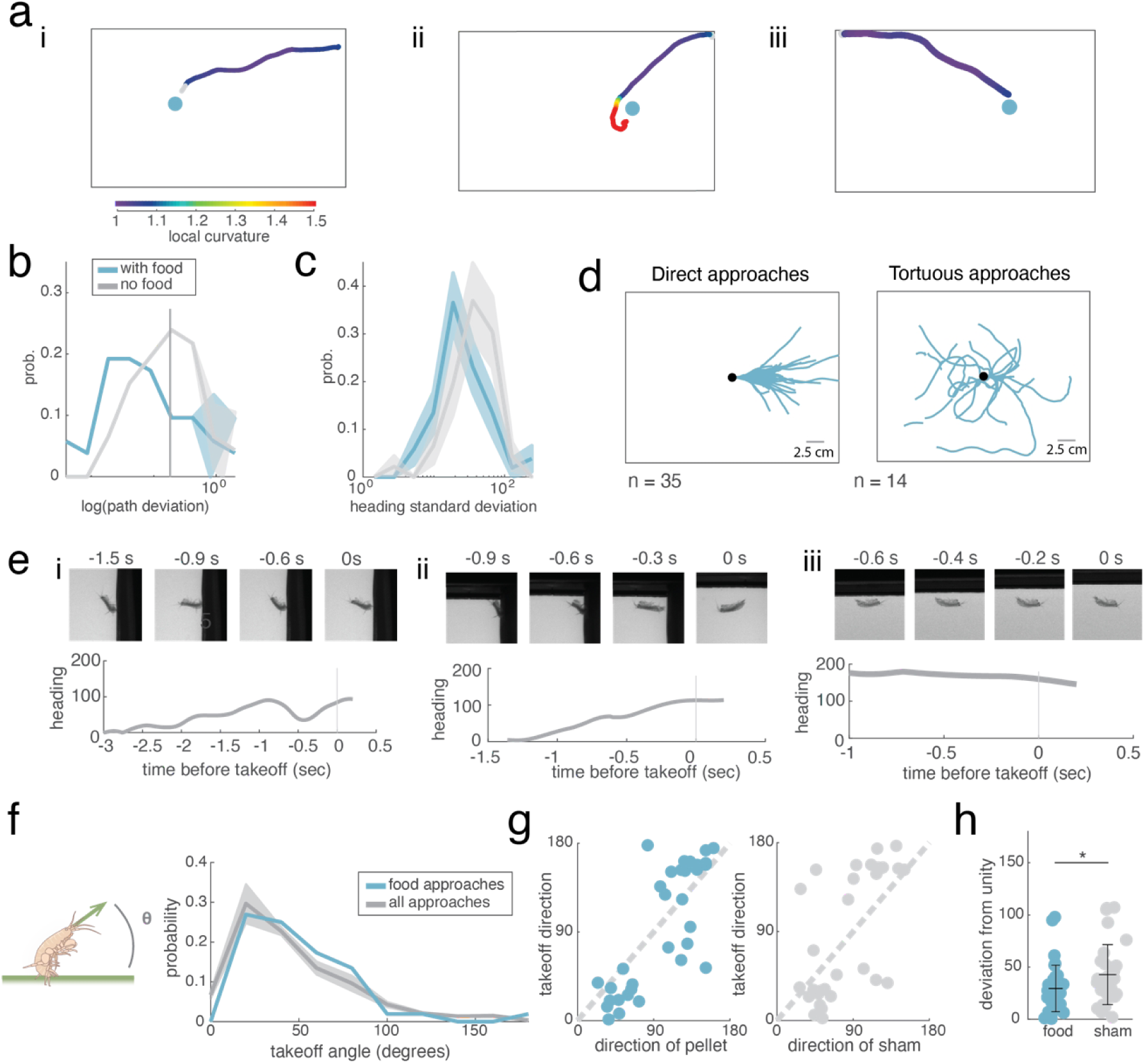
Intact animals utilize ballistic movements to approach food. a. Example approach to food for three intact animals, color coded by local path curvature (calculated as the arc-chord ratio over a window of 0.5 seconds). b. Distribution of path deviations (arc-chord ratio over the extent of a swim) for approaches to food pellet and sham pellet (KS test p = .0092) c. Distribution of heading standard deviations for approaches to food and sham pellet (KS test p = .0045) d. All direct (path deviation < 1.2) and indirect (path deviation > 1.2) food approaches aligned to final .2 second segment of approach. e. Example behaviors preceding approaches to food for the animals depicted in (a). Top: frames showing body angle positioning preceding takeoff. Bottom: measurement of animal’s heading relative to wall for period preceding takeoff. f. (right) illustration of takeoff angle calculation. (left) Distribution of takeoff angles for direct food approaches (blue) compared to the distribution of takeoff angles for all swims (gray), standardized to the animal’s initial heading direction. g. Left: Direction of food pellet at takeoff compared to the direction of swim takeoff for direct food approaches (R^2^ = 0.662, p=2.84 e-9). Right: Direction of sham pellet at takeoff compared to the direction of swim takeoff for direct sham approaches. (R^2^ = 0.376, p=.00024). Dashed line represents line of unity. h. Deviation from unity for plot of takeoff direction and pellet heading direction for direct approaches to food pellet and sham pellet (Wilcoxon Rank Sum, p = .035)

Where *θ̂*_-*i*_ represents the distribution produced by leaving out the *i*^th^ replicate, *θ̅* represents the mean of resampled distributions, and n represents the number of replicates.

## Results

### Food localization behavior in *Parhyale* requires the antennae

We first set out to characterize food search behavior of *Parhyale*. Intact *Parhyale* (Fig. 1a, N=49) were food-deprived for 9-17 days prior to experimentation. Each animal’s behavior was recorded in an experimental session and a control session, all of which occurred in the dark. In “experimental” trials, a food pellet and wax sham pellet were placed at two designated locations (Fig. 1b), which were shuffled between animals. Behavior was recorded for 120 minutes or until the animal found the food pellet, except for a subset of experiments (N=9), we continued imaging after food was found. Under these conditions, we found that animals spent extensive periods interacting with the food pellet (up to 1.5 hours, Supplemental Fig. 1c). In the other “control” session, no stimuli were provided, and behavior was recorded for 1 hour (Fig. 1d). Centroid position and orientation of the animal were tracked as described in Materials and Methods. For each animal, we then created an ethogram (Materials and Methods) to classify behavior at each time point into one of 6 categories: on wall, swimming, turning, stopped, on sham, or on food pellet (Fig. 1c,d).

We found that intact *Parhyale* readily found the food pellet and spent longer interacting with the food than with the sham pellet. 73.5% (35 of 49) *Parhyale* successfully found the food pellet (defined as spending >1 s on the target) compared to 46.9% (23 of 49) finding the sham (Fig. 1e). *Parhyale* also spent longer interacting with food than with the sham. To identify the longest recorded landing on the food or sham pellet, we capped the total interaction time at 120 seconds, the minimum duration recorded after food approach, to account for variability in recording protocols, and recorded *Parhyale* that did not approach the food as having a landing duration of zero. After doing so, we find that landings on food pellet are biased toward durations of more than 120 seconds, with a median capped duration of 113.4 seconds, compared to a median of 0.6 seconds for landings on the sham (Fig. 1f).

To gauge the significance of these behaviors for food localization, we performed the same analyses on behavior during sessions with no stimuli. We identified “pellet” and “sham” approaches for each animal as those that passed through the location of the food or sham pellet in the corresponding experimental trial. 38.8% of *Parhyale* in no food trials approached to location of the “food pellet”, while 36.7% approached the location of the “sham” (Fig. 1e). As we would predict, the median interaction time with both locations was 0 seconds (Fig. 1f). The presence of stimuli also promoted swimming behavior. During experimental trials, we found that *Parhyale* spent significantly more time engaged in active behaviors (other than wall-walking), than during no-stimulus sessions (Fig. 1g).

We next sought to gauge the contribution of the antennae to food localization. We removed both antennae from n=39 *Parhyale* and allowed them to recover during the food-deprivation period (see Materials and Methods). We imaged antennae-removed *Parhyale* using the same apparatus and protocol described above (Fig. 1h). Antenna-removed *Parhyale* were all imaged for the full 120 minutes in the presence of stimuli and 60 minutes with no stimuli, in pseudo-random order. We found that antenna removal severely disrupted food localization behaviors. Only 10.3% of full antenna removed animals found the food pellet, and they did not localize the food pellet more effectively than the sham (Fig. 1i, 15.4% of animals). Antenna-removed *Parhyale* did not spend significantly more time on the food pellet than the sham (Fig 1 j), with a median interaction time of 0 seconds each. These observations suggest that the antennae are required both the locate the food pellet and to discriminate the food pellet from the sham. Antennae-removed *Parhyale* also spent significantly less time swimming than intact animals (fig. 1k). Together these observations argue that *Parhyale* use their antennae to locate food and discriminate food from other objects, and that the antennae are also required for normal swimming behavior.

### *Parhyale* largely approach food using aimed ballistic movements

We next sought to examine food search behavior in greater detail. Detailed analysis of food search in animals navigating in low flow environments often demonstrates that these animals use many sequential sensory-driven turn and run decisions that collectively facilitate navigation up an odor gradient, bringing the animal closer to a food source (Gomez-Marin et al. 2011, Iino and Yoshida 2009). Instead, in intact *Parhyale*, we found that most food approaches were ballistic, with a single fixed trajectory for most of the period between leaving the wall and nearing the pellet (Fig. 2a, Supplemental Video 1).

To gauge whether such ballistic approaches were common, we extracted swim bouts preceding pellet approaches. We measured the directness of food approach trajectories as their path deviation, a ratio of the cumulative path length of the approach trajectory relative to the total displacement from start to finish, with a value of 1 indicating a perfectly straight path. Path deviations across all swims ranged from 1.008-298.95, with a mean of 1.85. The majority of food approaches (n = 35 of 49 approaches from 36 animals) had a low overall path deviation (<1.2, Fig. 2b), while control “approaches” (swims during control experiments that passed through the food pellet location in the corresponding experimental trial) were evenly divided between direct (n = 24 of 46) and indirect paths (b, Supplemental Fig. 2b). Food approaches were also less variable than control swims, having a lower standard deviation of heading (Fig. 2c).

To determine if *Parhyale* actively select the direction of these ballistic trajectories, we examined behavior immediately preceding take-off (Fig. 2e). We observed that some take-offs were preceded by a rearing behavior in which *Parhyale* dynamically adjusted their body angle relative to the wall (Fig. 2e left, center). Other animals gradually repositioned their body angle as they moved away from the wall. The distribution of heading angles adopted by *Parhyale* as they left the wall (relative to their heading direction along the wall) were biased toward shallow angles, peaking at 25° with and without food (Fig. 2f, Supplemental Fig.2a). To ask whether take-off angle was aimed at the food pellet, we plotted take-off angle as a function of direction of the pellet for all food-approach trajectories (Fig. 2g). We found that these were well-correlated (R^2^ = 0.66, p = 2.84e-9) and were closer to unity for pellet approaches than for sham approaches (R^2^ =.38, p = .035, Fig. 2g,h, Supplemental Fig.2c-e). We conclude that most pellet approaches in *Parhyale* represent aimed ballistic movements in which the angle of approach is chosen while still at the wall.

### Aiming and course maintenance require the antennae

Given that removal of both antennae severely disrupted food pellet finding, we wondered if antennal removal also disrupts aiming and ballistic approaches. Examining trajectories from antenna-removed animals revealed a profound deficit in the ability to maintain a straight heading (Fig. 3 a,b). While intact animals were able to swim with consistent headings (Fig. 3a), full antenna removed animals exhibited meandering and tortuous paths, despite similar average movement speeds (Fig. 3b, Supplemental Fig. 3). Over each continuous period of swimming and turning (excluding stops, and time on walls, pellet, or sham) antenna removed animals had higher average angular velocities (Fig. 3c) and exhibited increased variability in their heading direction and angular velocity (Fig. 3d). *Parhyale* with both antennae removed also showed a deficit in aiming behavior, as antenna removed animals were more likely to take off from the wall at larger angles (Fig. 3e), suggesting that the antennae are required both for selecting an appropriate take-off angle, and to maintain a straight heading and course during swims. These data indicate that antennae play a key role in both chemosensation and control of orientation during locomotion in *Parhyale*.

**Figure 3.**
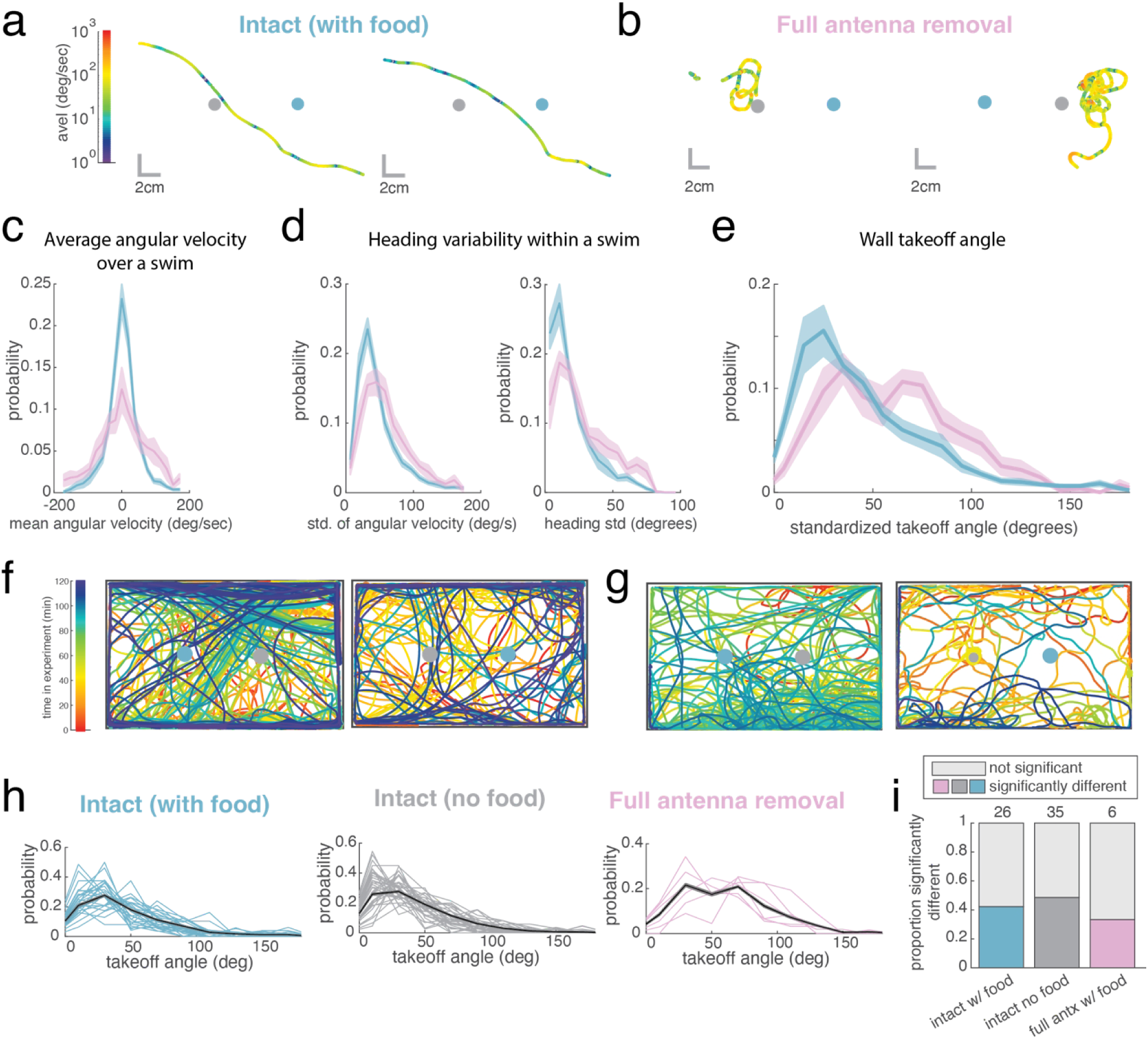
Antennae are necessary for *Parhyale* to maintain a stable heading. a. Example swim traces from an intact animal, color coded by instantaneous velocity. b. Example swim traces from a full antenna removed animal, color coded by instantaneous velocity. c. Distribution of angular velocities during swimming for intact and full antenna removed animals (KS test p= 7.25 e-6). d. Left: standard deviation of angular velocity (measured by animal’s eye to body orientation) over all swims for intact full antenna removed animals (KS test p = 0.0028). Right: standard deviation of heading direction (KS test p = 2.87 e-5) e. Distribution of takeoff angles for intact and full antenna removals (KS test p = 3.3 e-18), standardized to the initial direction the animal was facing. f. Example swimming from intact animals, color coded by time during experimental trial. g. Example swimming from full antenna removed animals, color coded by time during experimental trial. h. Distribution of standardized takeoff angles for animals that swim robustly (>25 takeoff events), with population distribution (black) for intact experiments with food (left), without food (middle), and full antenna removed with food (right). i. Proportion of individuals that exhibit significantly different distributions of takeoffs from the population average (defined as KS test p<.05) across intact, control, and first antenna removed groups

We observed that intact animals often followed similar trajectories over successive swims through the arena (Fig. 3f), and that consequently, individual animals often exhibit biased distributions of takeoff angles relative to the population mean (Fig. 3h). However, in full antenna removed animals, takeoff headings were more likely to align with the population average (fig. 3g). To determine if this observation was consistent across animals, we extracted takeoff distributions for animals that swam robustly (> 25 swims) and isolated their takeoff distributions (Fig. 3h). We then quantified the proportion of individuals that significantly deviated from the population average distribution (by KS test) and found that about 42% of intact animals exhibited biased patterns of takeoff angles, compared to about 33% of full antenna removed animals (Fig. 3i).

### First and second antennae redundantly contribute to food finding and trajectory control

Given the pronounced effects of removing both antennae on behavior, we wondered whether the two antennal systems play distinct roles. Because crustaceans have previously been shown to use either their first antennae (in decapods, Steullet et al. 2002; Horner et al. 2004) or their second antennae (in terrestrial isopods, Kenning and Harzsch 2013; Seelinger 1983) as their primary chemosensory organ, we selectively removed either the first or second antennae bilaterally to investigate their individual contributions to food search behaviors (Fig. 4a,b). We imaged food search behavior in these partial antennal ablation conditions using the same paradigm as for full antennal ablations (Fig. 4 a,b). Unlike full antenna removals, both first- and second- antennae ablated animals were readily able to locate food (Fig. 4c), and were only slightly less successful than intact animals with 54.1% of first antenna removed animals and 56.8% of second antenna removed animals locating the food pellet.

**Figure 4.**
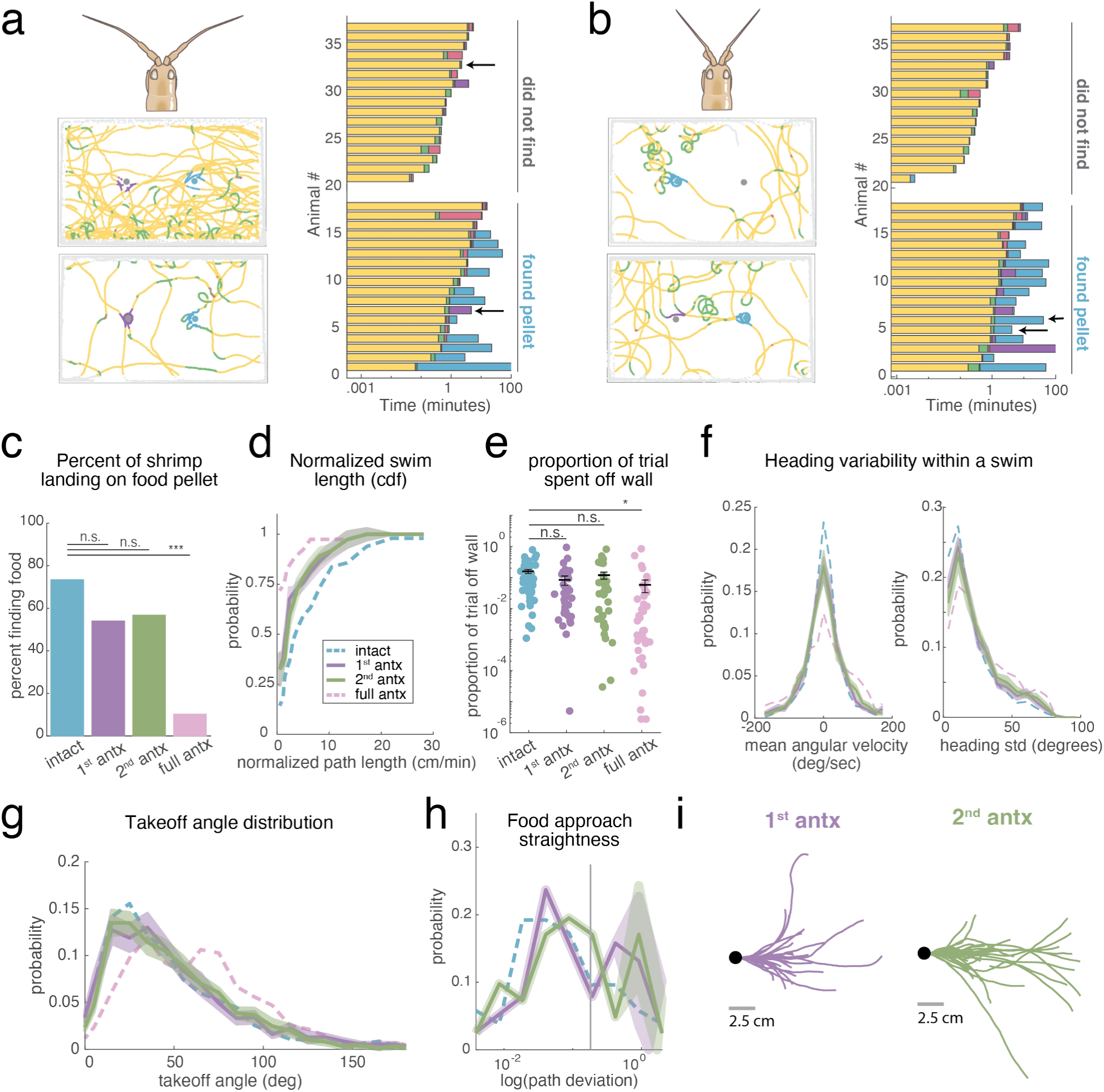
First and second antennae redundantly drive food search. a. Top: illustration of first antenna removal. Left: examples of locomotion in first antenna removed animals in the presence of food, color coded by behavioral state (swim, turn, stop, sham, pellet, wall walking); (right) ethogram of active behaviors (excluding wall walking) in the presence of food for all first antenna removal experiments (N = 37), arrows indicating experiments used as examples. b. Top: illustration of second antenna removal. Left: examples of locomotion in second antenna removed animals in the presence of food, color coded by behavioral state (swim, turn, stop, sham, pellet, wall walking); (right) ethogram of active behaviors (excluding wall walking) in the presence of food for all second antenna removal experiments (N = 37), arrows indicating experiments used as examples. c. Percent of animals finding food in first antenna removal experiments (Chi Sq. p= 0.0614 compared to control) and second antenna removal (Chi Sq. p= 0.1045 compared to control). d. CDF of normalized swim length (total path length standardized to length of experiment) □ jackknife standard deviation for first and second antenna removed animals (KS test p = .8631). Intact distribution (1^st^ antx KS test p = .0084; 2^nd^ antx KS test p =.1101) and full antenna removal distribution (1^st^ antx KS test p = 2.12 e-5; 2^nd^ antx KS test p = .0002) included for reference. e. Proportion of experiment spent engaged in swimming (non wall-walking) behaviors for 1^st^ antenna removals (ANOVA p = .1857 relative to intact, p = 0.9026 relative to full antx) and 2^nd^ antenna removals (ANOVA p = 0.7224 relative to intact, p = 0.3805 relative to full antx). f. Left: distribution of average angular velocities over individual swims, for 1^st^ and 2^nd^ antenna removed animals (KS test p =3.81 e-5). Intact (KS test vs 1^st^ antx p = 2.17 e-7, 2^nd^ antx p = 1.24 e-8) and full antenna removal (KS test vs 1^st^ antx p = 1.3 e-5, 2^nd^ antx p = .667) traces included for comparison. Right: distribution of heading standard deviation over individual swims, for 1^st^ and 2^nd^ antenna removed animals (KS test p = 2.67 e-6). Intact (KS test vs 1^st^ antx p = 2.64 e-27, 2^nd^ antx p =2.67e-11) and full antenna removal (KS test vs 1^st^ antx p = 5.86e-14, 2^nd^ antx p =0.01) traces included for comparison. g. Distribution of takeoff angles, standardized to initial heading direction ± jackknife standard deviation for all swims for 1^st^ and 2^nd^ antenna removed animals (KS test p = 0.505). Intact (KS test vs 1^st^ antx p = 0.0066, 2^nd^ antx p = 0.0228) and full antenna removal (KS test vs 1^st^ antx p = 1.6 e-11, 2^nd^ antx p = 4.67 e-10) traces included for comparison. h. Distribution of overall path deviation for food approaches in 1^st^ and 2^nd^ antenna removed animals (KS test p =.3805). i. Direct (path deviation < threshold) food approaches for 1^st^ and 2^nd^ antenna removed animals, aligned to path at final 0.2 seconds of approach.

Beyond their capacity for finding food, the locomotion of first- and second- antenna removed animals was consistently intermediate between that of the intact and full antenna removed *Parhyale*. First and second antenna removed animals had slightly reduced overall path lengths compared to intact animals despite similar average velocities (Fig 4d, supplemental Fig. 4d) and spent a similar proportion of the trial swimming compared to intact animals (Fig. 4e). As with the full antennal ablations, both the first and second antenna removed animals exhibited moderate increases in both the average angular velocity and standard deviation of heading across swim trajectories, but only a slight increase in average path curvature (Fig. 4f, Supplemental Fig. 4a-b). However, both first and second antenna removed animals consistently took off from the wall at similarly shallow angles to intact animals (Fig. 4 g), suggesting that both first and second antennae removed animals retain the ability to estimate and control their orientation. Supporting that hypothesis, we observed low-curvature food approaches in both antenna removal conditions (Fig. 4h-i). Together these observations argue that the first and second antennae of *Parhyale* play largely redundant roles in the control of orientation during locomotion.

### Antennae contribute to food modulated turns and stops

While most food approaches were ballistic, we often observed turns close to the pellet facilitating landing (e.g. Fig. 2aii). Additionally, a minority of approaches were more tortuous, and featured multiple turns or stops near the food pellet (Fig. 5 a,b). Plotting the location of turns and stops across our recordings from intact *Parhyale* (Fig. 5c) revealed that animals turned preferentially in the vicinity of the food pellet, rather than the sham pellet (Fig. 5d-i). The median turn distance from the pellet (3.75 cm) was significantly lower than the median turn distance from the sham (4.72 cm, Fig 5d-iii). The joint distribution of turn angular velocity and turn distance from the food pellet shows that high angular velocity turns occur preferentially near the food pellet, indicating that they may be characteristic of food search (Fig. 5d-iv). No such difference in the distribution of turn and stop locations was observed during session without stimuli, arguing that these behaviors reflect a sensory response to the food pellet (Fig. 5b: median distance from “pellet”= 4.27cm; median distance from “sham”= 4.24 cm).

**Figure 5.**
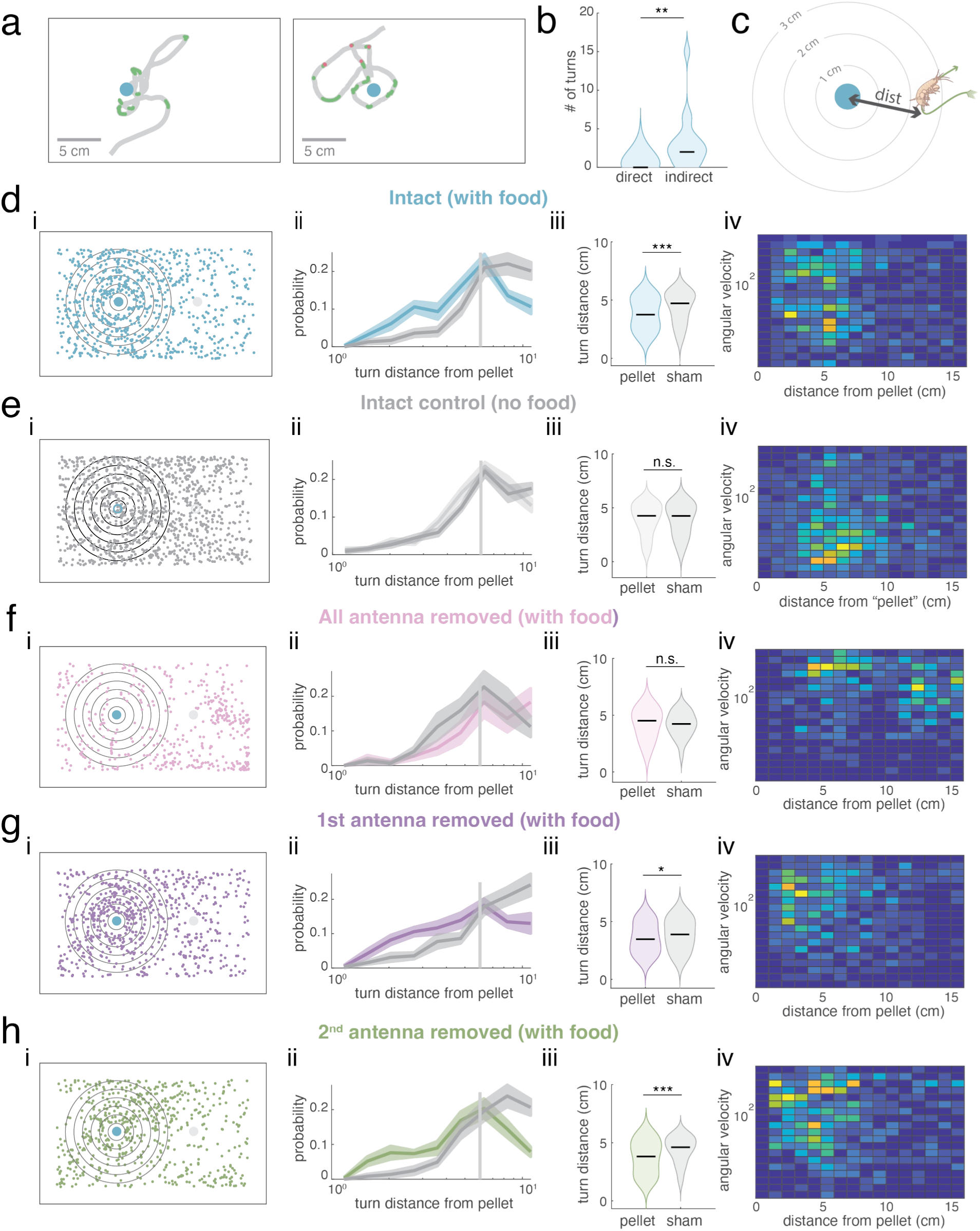
Turning phenotype suggests first- and second- antenna removed animals remain responsive to odor presence and proximity. a. Example indirect food approaches with stops (pink) and turns (green) highlighted. b. Number of turns during food approach swim for low and high path deviation approaches. Black line indicates median turn count, (rank sum, p = .0002) c. Illustration of turn distance calculation. d. Turn features for intact animals with food. (i) Turn location (center of mass) plotted in arena coordinates aligned to pellet location (X-axis reversed in trials where the food pellet was placed on the right). Large dots indicate location of food pellet (blue) and sham pellet (grey). Grey outlines indicate 1 cm increments from food pellet. (ii) Probability of observing turn as a function of distance from food or sham pellet (KS test p = 1.01 e-16) (iii) Within 6cm radius, average turn distance relative to food pellet and sham pellet (rank sum p =.00015), black lines indicate median. (iv) Joint distribution of turn angular velocity and distance from pellet. e. Turn features for intact animals without food. (i) Turn location (center of mass) plotted in arena coordinates aligned to “pellet” location in corresponding with food trial. Large dots indicate location of “food”(blue) and “sham” (grey). Grey outlines indicate 1 cm increments from “food” location. (ii) Probability of observing turn as a function of distance from “food” or “sham” location (KS test p = 8.35 e-5) (iii) Within 6cm radius, average turn distance relative to food pellet and sham pellet (rank sum p = .7559), black lines indicate median. (iv) Joint distribution of turn angular velocity and distance from pellet. f. Turn features for full antenna removed animals with food. (i) Turn location, as in (a) (ii) Probability of observing turn as a function of distance from food or sham pellet (KS test p = 1.17 e-8) (iii) Within 6cm radius, average turn distance relative to food pellet and sham pellet (rank sum p = 0.2233), black lines indicate median. (iv) Joint distribution of turn angular velocity and distance from pellet. g. Turn features for first antenna removed animals with food. (i) Turn location, as in (a) (ii) Probability of observing turn as a function of distance from food or sham pellet (KS test p = 1.26e-15) (iii) Within 6cm radius, average turn distance relative to food pellet and sham pellet (rank sum p = .0346), black lines indicate median. (iv) Joint distribution of turn angular velocity and distance from pellet. h. Turn features for full antenna removed animals with food. (i) Turn location, as in (a) (ii) Probability of observing turn as a function of distance from food or sham pellet (KS test p = 6.16 e-13) (iii) Within 6cm radius, average turn distance relative to food pellet and sham pellet (rank sum p = .0003), black lines indicate median. (iv) Joint distribution of turn angular velocity and distance from pellet.

If turns and stops near the food pellet represent a sensory response, we would expect them to depend on the antennae. Consistent with this hypothesis, removing both antennae completely abolished food-specific turning (Fig. 5c-i) and increased the median distance of turns from the pellet (Fig. 5 c-iii, pellet median = 4.5 cm, sham median = 4.2 cm). In *Parhyale* with either the first or second antennae removed, turns were more common near the food pellet than the sham pellet, as in intact animals. In first antenna removals, the median turn distance was 3.48 cm from the food and 3.88 cm from the sham pellet (Fig. 5 d-iii,). Likewise, in second antenna removals, turns were significantly closer to the food (median distance = 3.81 cm) than the sham pellet (median = 4.6 cm, Fig. 5 e- iii). In both groups, the joint distribution of angular velocity and turn distance shows a cluster of high angular velocity turns occurring near the pellet (Fig. 5 d,e-iv). This suggests that either set of antennae is sufficient for *Parhyale* to detect the presence of and proximity to a food pellet. However, unlike intact animals, first and second antenna removed animals did not show an increased probability of stopping near food (Supplemental Fig. 5), indicating some impairment to food-related behaviors associated with single antenna loss. Together these observations argue that both antennae contribute to food sensing— likely through chemosensation— allowing for turns in proximity to the food pellet.

### Antenna removal alters turn structure

Finally, to determine if antenna removal disrupted heading stability during reorientations, we examined the detailed structure of turns. Turns were identified based on high local curvature (as described in ethogram section above, Fig. 6a) In intact animals, turns covered a similar total distance with and without food present (Fig. 6 b, c), but turns in the presence of food were more likely to have a high angular velocity (Fig. 6d). Antenna-removed *Parhyale* spent a greater proportion of their swimming engaged in turning (average of 26% of total swimming) and stopping (21.2% of swimming) compared to intact animals (10.2% turning and 8% stopping).

**Figure 6.**
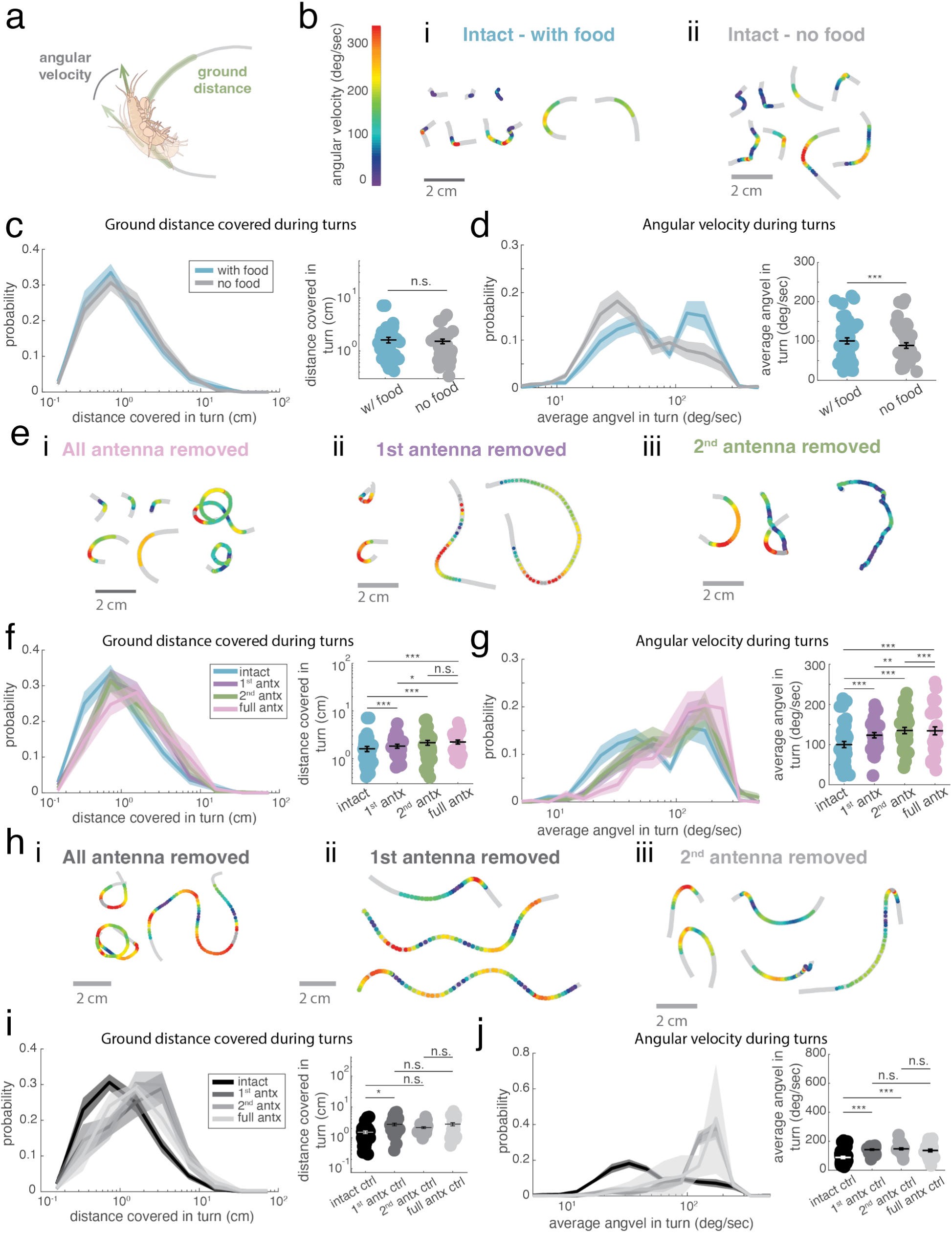
Antenna removal destabilizes turn structure. a. Illustration of turn calculations demonstrating angular velocity and ground distance covered. b. (i) Examples of turns in intact animals in the presence of food, color coded by instantaneous angular velocity (degrees/sec). Grey trace shows overall trajectory for 0.2 seconds preceding and following turn. (ii) Examples of turns in intact animals without food, color coded by instantaneous angular velocity (degrees/sec). c. (left) Distribution of ground distance covered during turns for intact animals with and without food (KS test p =0.063). (right) Average distance covered during turn, for intact animals with and without food (rank sum p = .1237). d. (left) Distributions of average angular velocity during turns for intact animals with and without food (KS test p =5.73 e-10) (right) Average turn angular velocities during turn, for intact animals with and without food (rank sum p = 8.58 e-12). e. Example turns from full antenna removed (left), 1^st^ antenna removed (middle), and 2^nd^ antenna removed animals (right) in the presence of food, color coded by instantaneous angular velocity (degrees/sec). Grey trace shows overall trajectory for .2 seconds preceding and following turn. f. Left: Distribution of distances covered in turns during food present experiments for antenna removed animals relative to intact animals in with food experiments. Full, 1^st^ and 2^nd^ antenna removed animals, compared to intact turn distance distributions (KS test full antx p = 2.9 e-8, 1^st^ antx p = 6.35 e-5, 2^nd^ antx p = 2.05 e-5). 1^st^ and 2^nd^ antenna removals compared to full antx distributions (KS test 1^st^ antx p = 0.0725, 2^nd^ antx p =0.1281) included as a control. Right: Average ground distance covered during turns in experiments with food, for all animals. Full antenna removed animals relative to intact (rank sum p = 1.96 e-9), 1^st^ and 2^nd^ antx compared to intact animals (rank sum 1^st^ antx p = 1.58e-5, 2^nd^ antx p =2.01 e-6) and full antx animals (1^st^ antx p = 0.012, 2^nd^ antx p = 0.0766 relative to full antx). g. Left: Distribution of turn angular velocities during food present experiments for all animals. Comparing full antx animals to intact (KS test relative to intact p = 7.6 e-14). 1^st^ and 2^nd^ antenna removed animals (KS test p = 0.426), compared to intact distribution (KS test 1^st^ antx p = 1.29 e-8, 2^nd^ antx p = 7.84 e-6) and full antenna removed distributions (KS test 1^st^ antx p = 0.0245, 2^nd^ antx p = 0.0011) Right: Average turn angular velocity during experiments with food, comparing full antenna removals to intact animals (rank sum p = 6.65 e-15 relative to intact); 1^st^ and 2^nd^ antenna removals to turns in intact animals (rank sum 1^st^ antx p = 1.66e-8, 2^nd^ antx p = 1.29 e-6); and to turns in full antenna removed animals (rank sum 1^st^ antx p = 0.0019, 2^nd^ antx p = 0.0005). h. Example turns from full antenna removed (left), 1^st^ antenna removed (center) and 2^nd^ antenna removed animals (right) in no food trials, shown as in (a). i. Left: Distribution of distances covered in turns during no food experiments for 1^st^ and 2^nd^ antenna removed animals, with intact distribution (KS test 1^st^ antx p = 7.1 e-12, 2^nd^ antx p = 2.19 e-19) and full antenna removed distributions (KS test 1^st^ antx p = .1361, 2^nd^ antx p = .0521) included as a control. Right: Average ground distance covered during turns in experiments without food, comparing 1^st^ and 2^nd^ antenna removals to turns in intact animals (rank sum 1^st^ antx p = 4.06 e-11, 2^nd^ antx p =3.45 e-20) and to turns in full antenna removed animals (rank sum 1^st^ antx p = .6494, 2^nd^ antx p = .1298). j. Left: Distribution of turn angular velocities during no food experiments for 1^st^ and 2^nd^ antenna removed animals (KS test p = .5626), with intact distribution (KS test 1^st^ antx p = 3.33 e-52, 2^nd^ antx p = 2.81 e-68) and full antenna removed distributions (KS test 1^st^ antx p = .0786, 2^nd^ antx p =.0136) included for comparison. Right: Average turn angular velocity during experiments without food, comparing 1^st^ and 2^nd^ antenna removals to turns in intact animals (rank sum 1^st^ antx p = 8.54 e-49, 2^nd^ antx p = 9.24 e-56) and to turns in full antenna removed animals (rank sum 1^st^ antx p = .7873, 2^nd^ antx p = .4782).

Turn structure was also changed by antenna removal. Unlike turns in intact animals, which were generally brief reorientations, antenna removed animals exhibited long, meandering turns (Fig 6e). In antenna removed animals, turns covered a greater total ground distance (Fig. 6f) and had higher angular velocity (Fig. 6g) than turns from intact animals, consistent with observed increase in path instability. First and second antenna removed animals also exhibited increased angular velocity and total distance covered during turns, which once again was intermediate between the full antenna removal and the intact animals (Fig.6 f-g). Notably, however, turns produced during control experiments (in the absence of food) by first and second antenna removed animals closely resemble those of full antenna removals (Fig. 6 h-j), and were biased towards longer ground distances and higher angular velocities, resembling the turning behaviors of full antenna removed controls. While we cannot determine to what extent changes to turn structure in antenna removed animals are a byproduct of a sensory defect (i.e., an inability to detect self-motion) versus an active strategy employed by animals to compensate for the sensory deficit, this observation suggests that animals missing only the first or second antennae may be capable of controlling their turning behaviors when food is present to facilitate food localization.

## Discussion

In this study, we have quantitatively described the food search behaviors of *Parhyale* hawaiensis. We find that these animals make ballistic, targeted movements when approaching food, which depend on the presence of antennae. Additionally, we demonstrate that antennae are necessary for orientation and maintenance of heading during swimming. Finally, we show that *Parhyale* exhibits similar behavioral responses to first or second antenna removal, suggesting that these animals have the capacity to utilize either set of antennae in their food search behavior.

### *Parhyale* perform a novel food approach behavior

*Parhyale* most commonly approached food using ballistic, highly oriented swims from the arena wall (Fig. 2), a distinct behavior from the gradient search strategies of other arthropods (Gomez-Marin et al. 2011; Gomez-Marin and Louis 2014). Because both orientation from the wall and maintenance of heading during swimming were disrupted by antennae removal, we can conclude that these features of swimming are actively controlled based on sensory feedback. Directed runs are a common component of olfactory navigation strategies. In larval *Drosophila*, which navigate in a low-flow environment like that presented to *Parhyale*, “weathervaning” runs are directed by active head casting behavior (Gershow et al. 2012; Gomez-Marin et al. 2011). In more dynamic environments, orientation toward wind (anemotaxis) is a commonly observed navigation strategy (Matheson et al. 2021; Kennedy and Marsh 1974; Stupski and van Breugel 2024).

Following these two models, we hypothesize that *Parhyale*’s ballistic runs are targeted either by measurement of the odor gradient immediately prior to takeoff, or by estimation of the food location based on prior exploration. In insects, maintenance of a straight course during olfactory navigation is associated with short term representations of a goal direction relative to global environmental cues in the Central Complex (Kathman et al. 2026; Lanz et al. 2025; Westeinde et al. 2024; Stone et al. 2017; Mussells Pires et al. 2024). *Parhyale* possess a clearly delineated Central Complex consisting of a protocerebral bridge, a central body (with upper and lower components homologous to the fan-shaped body and ellipsoid body in *Drosophila*), and lateral accessory lobes (Wittfoth et al. 2019). As such, a potential substrate for directional representations exists in these animals. Future experiments testing the efficacy of *Parhyale*’s food search behavior in the presence of a strong directional cue (such as flow), or in the absence of directional cues (such as in a circular arena), would help to elucidate if ballistic approaches depend on directional information.

Why might *Parhyale* preferentially utilize ballistic approach over gradient search? *Parhyale* are known to inhabit intertidal regions and mangrove swamps (reviewed Paris et al. 2022) regions in which tidal currents greatly exceeding the animal’s own locomotor speed can create complex and dynamic flow regimes (Wolanski 1992). Ballistic launches from a fixed substrate, akin to a swimmer kicking off of a wall, might allow an animal to maximize the distance it can cover in a short period of time, limiting the risk of being driven off course by strong flow. Further examination of *Parhyale*’s locomotion across flow conditions and substrates may help to clarify if and how environmental stability biases locomotor decisions.

### Why does antenna removal result in heading instability?

Following lesions of all antennae, *Parhyale* exhibited increased heading instability during swimming, and reduced consistency in their wall takeoff headings. We also find that turns during food search become longer and sharper after antenna removal. Insects detect airflow via a chordotonal organ known as Jonston’s organ, which senses deflection of the third antennal segment (Kamikouchi et al. 2009; Göpfert and Robert 2002) and contributes to detection of self-motion during flight (Budick et al. 2007; Fuller et al. 2014; van Breugel et al. 2022). While the mechanisms of flow sensing are less clear in crustaceans, crustacean antennae possess abundant mechanosensors. These include a chordotonal organ at the base of the first antennal flagellum resembling the insect Jonston’s Organ, and mechanosensory sensilla on the first and second antennae which respond to displacement due to water flow (reviewed Watling and Thiel 2013; Harzsch and Krieger 2018), making self-motion detection via the antennae plausible in crustaceans. In some crustaceans, an additional organ of equilibrium in the basal segment of the first antennae, the statocyst, detects self-motion to drive orienting reflexes (Leedham et al. 2026, Davis 1968). If *Parhyale* also possess these organs in the first antennae, this might explain the moderate increase in average path curvature we observe following first, but not second antenna ablation (Supplemental Fig. 4 a,b). However, the similarity of orientation phenotypes across antennal manipulations suggests a sensory control of locomotion that extends beyond any single receptor system.

*Parhyale* exhibit dynamic locomotor behaviors and may prove to be a valuable system for studying sensory control of locomotion. Although outside of the scope of this paper, across our experiments we observe that *Parhyale* adopt a range of body postures during locomotion (Supplemental Video 1) and will undergo dynamic reorientations during swimming. Selective ablation of chemosensory neurons (as in Steullet et al. 2001) could establish if the impairment to orientation presented here arises solely due to deficits to chemosensation. Further manipulation of mechanosensation by stabilizing antennal joints (to disrupt the function of chordotonal organs, as in Álvarez-Salvado et al. 2018) or of statocysts by introducing magnetic grains to be incorporated into the statocyst organ (as in Ozeki et al. 1978) may help to parse how these separate sensory streams contribute to locomotion.

### Behavioral redundancy of 1^st^ and 2^nd^ antennae

In our experimental paradigm, we found significant redundancy in the contributions of first and second antennae to behavior. In *Parhyale*, like other crustaceans, chemosensory signals at the antennae are processed in three different channels. Unimodal chemosensory sensilla on the first antennae house olfactory sensory neurons (OSNs), which project to glomeruli in the olfactory lobes, while bimodal chemo-and mechanosensory sensilla on both antennae contain distributed chemoreceptive neurons (dCRNs) that project to antenna specific neuropils (Kümmerlen et al .2025.; Wittfoth et al. 2019). Prior work has largely assumed that the chemosensory capacity of the second antenna is utilized for contact chemosensation (i.e. taste), while the unimodal and bimodal chemoreceptors of the first antenna mediate odor identity determination and spatiotemporal discrimination necessary for navigation, respectively (Schmidt 2007, Mellon 2007, Mellon 2012). However, based on the behavioral evidence of this paper and others (Dunham et al. 1997; Voigt and Atema 1992; Kenning and Harzsch 2013; Seelinger 1983), we propose that the distributed chemosensory systems of the first and second antenna fulfill similar functions in several crustacean species.

Why might *Parhyale* have developed such a dramatic redundancy in its chemosensory system? One explanation is that comparisons between antennae can be used to extract features of odor structure such as concentration gradients over space (Lockey and Willis 2015). Prior work in crayfish (Kraus-Epley and Moore 2002), suggests that comparisons across the antennae are an important part of crustacean chemosensory navigation strategy. Increasing antenna number might therefore provide redundancy to the system in case of antennal injury. Alternatively, antennal redundancy might compensate for active sensing behaviors (flicking of the first antennae) that are akin to sniffing in crustaceans (Mead and Koehl 2000; Reidenbach et al. 2008). A separate chemosensory system on the second antenna might therefore enable steady tracking of odor and background flow while the first antennae are involved in active sampling.

## Conclusion

*Parhyale* exhibit a unique, ballistic food approach strategy. While it remains unknown how this strategy is generated in the brain, *Parhyale* possess the conserved neural substrate for that facilitates orientation toward goals during navigation in insects, which may be utilized to drive this behavior. We additionally observe that antenna removal increases heading instability in swimming *Parhyale*, most likely due to a deficit in mechanosensation. Finally, we find that *Parhyale* exhibit robust food localization following the removal of either first or second antennae, with similar behavioral phenotypes. This is a novel demonstration of a crustacean that appears to flexibly utilize first or second antenna mediated chemosensation for food localization and orientation during locomotion. In this work, we establish a new experimental paradigm that can be used to investigate sensory and locomotor neurobiology in an accessible crustacean model.

## Acknowledgements

We thank Nipam Patel and Heather Bruce for providing Parhyale. We thank Quentin Gaudry and Aaron True for advice on building our recirculating behavioral arena. We thank Marie Suver for guidance on DeepLabCut and Yasha Zaman and Doyeon Sung for assistance with training dataset annotation. This work was supported by NSF 1559641 Odor2Action and NIH R34DA059500.

---

**Supplemental Figure 1.**
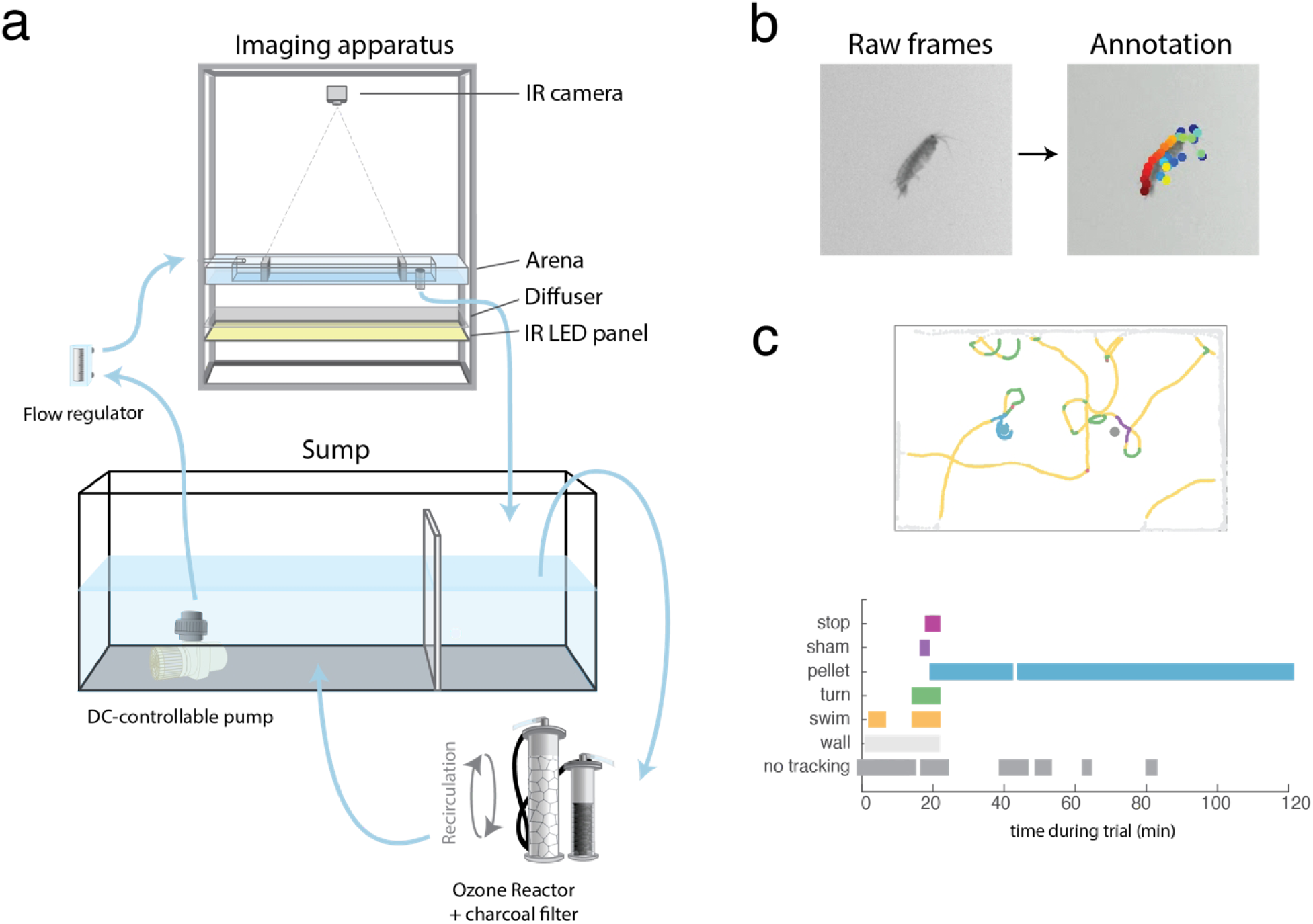
Methods for monitoring locomotor behavior in *Parhyale*. a. Schematic of flow apparatus. b. Example of isolated frame from experimental video, and the same frame annotated using DeepLabCut. c. (top) Example trace for an intact experimental trial that ran the full 120 minute duration. (bottom) illustration of ethogram phase at across trial duration.

**Supplemental Figure 2.**
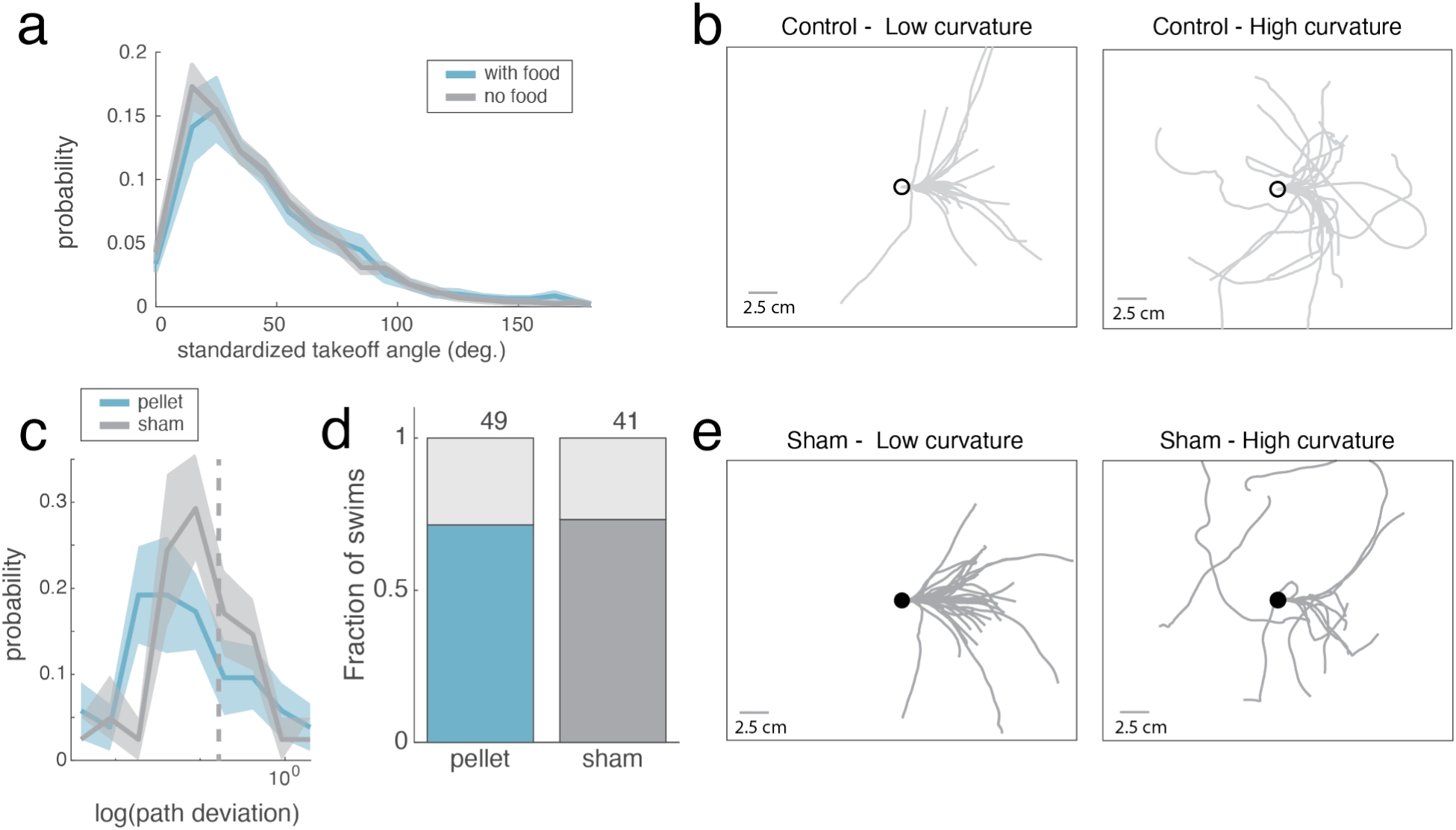
Orientation during control and sham pellet approaches. a. Distribution of takeoff angles for intact animals swimming with and without food (KS test p = 0.006). b. All high and low path deviation “food approaches” in control trials, oriented to the final 0.2 second of approach. c. Distribution of path deviations for approaches to food and sham (KS test p = 0.0412). d. Fraction of food and sham approaches that are direct and indirect. e. High and low path deviation sham approaches, oriented to the final 0.2 seconds of approach.

**Supplemental Figure 3.**
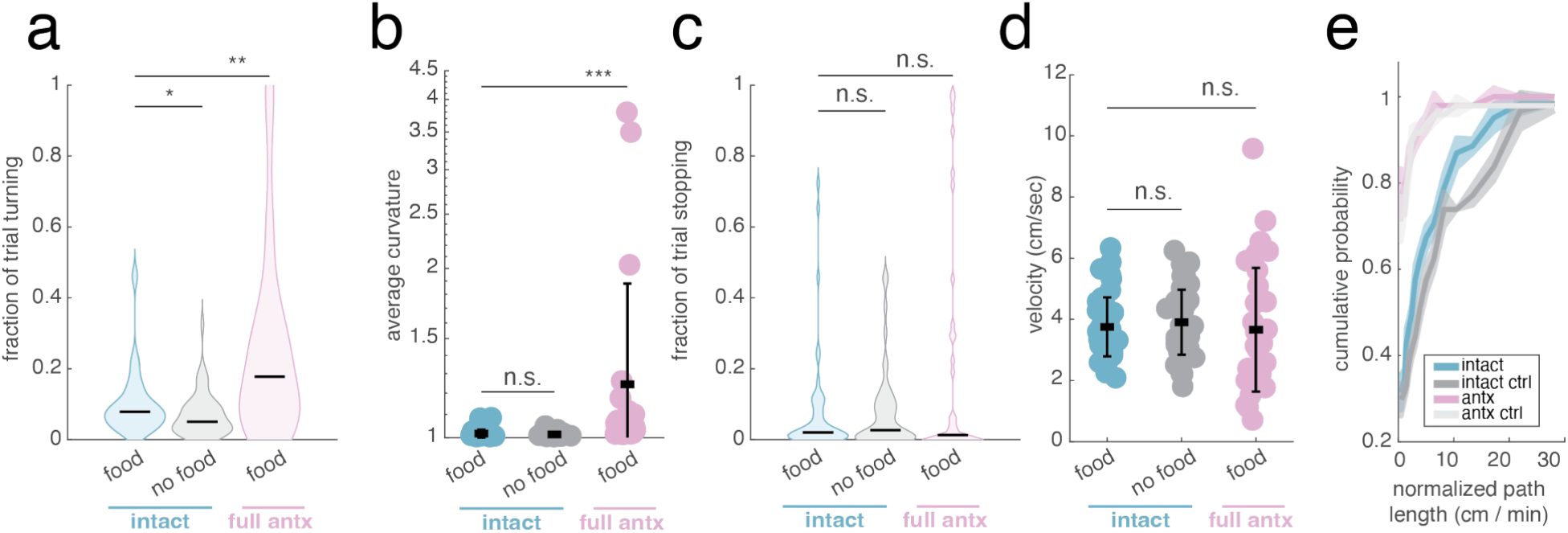
Antenna removal alters curvature, but not velocity related metrics. a. Fraction of active locomotion (ethogram swimming, turning, and stopping phase) spent turning for intact animals (with food vs control, rank sum p = .0388), and full antenna removed animals with food (compared to intact, p = .0037) b. Average path curvature during ethogram swimming and turning phases. Intact with vs without food (rank sum p = .0733), and intact vs full antenna removed with food (p = 3.32 e −10). c. Fraction of active locomotion (ethogram swimming, turning, and stopping phase) spent stopped for intact animals (with food vs control, rank sum p = 0.74), and full antenna removed animals with food (compared to intact, p = 0.899) d. Average velocity during ethogram swimming and turning phases. Intact with vs without food (rank sum p = .4453), and intact vs full antenna removed with food (p = 0.2217). e. Cumulative distribution of total path length (normalized by experiment duration). (KS test, intact experimental vs control p = 0.3459, antenna removed experimental vs control = 0.8178, intact experimental vs antenna removed experimental p = 2.083 e-8)

**Supplemental Figure 4.**
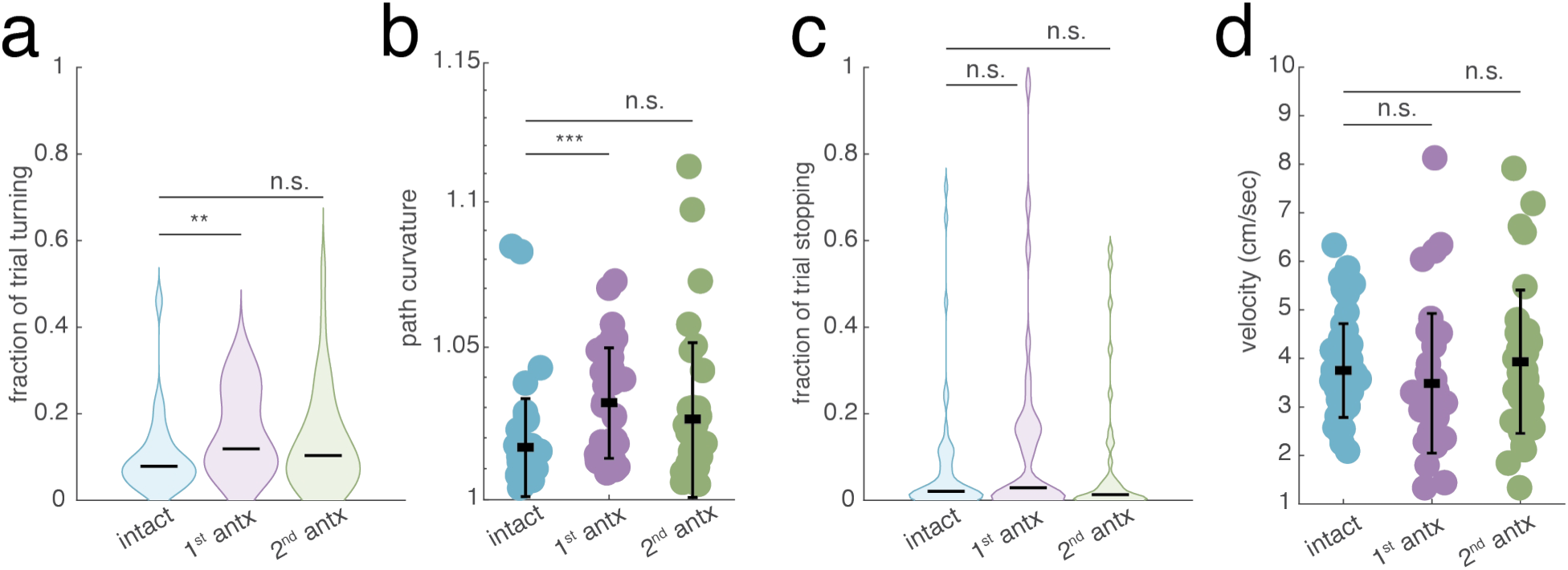
Gross locomotor features of single antenna removal conditions. a. Fraction of active locomotion (ethogram swimming, turning, and stopping phase) spent turning for 1^st^ antenna removed animals (vs intact, rank sum p = .0024), and for 2^nd^ antenna removed animals (vs intact, rank sum p = 0.2172) b. Average path curvature during ethogram swimming and turning phases1^st^ antenna removed animals (vs intact, rank sum p = 4.41 e-6), and for 2^nd^ antenna removed animals (vs intact, rank sum p = 0.071) c. Fraction of active locomotion (ethogram swimming, turning, and stopping phase) spent stopped for 1^st^ antenna removed animals (vs intact, rank sum p = .3082), and for 2^nd^ antenna removed animals (vs intact, rank sum p = 0.488) d. Average velocity during ethogram swimming and turning phases for 1^st^ antenna removed animals (vs intact, rank sum p = .0629), and for 2^nd^ antenna removed animals (vs intact, rank sum p = 0.6114)

**Supplemental Figure 5.**
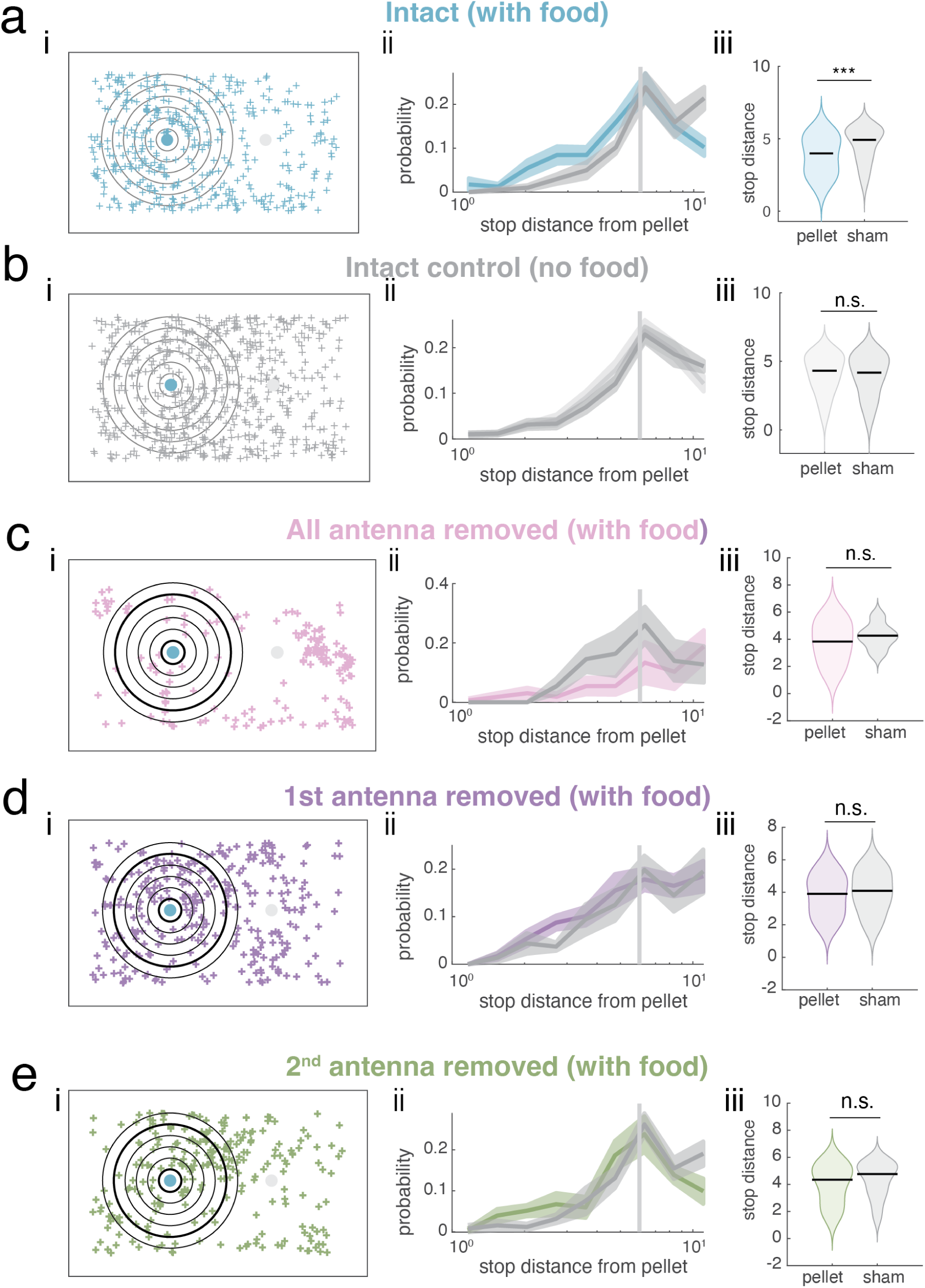
Antenna removed animals are less likely to stop near food. a. Stop features for intact animals with food. (i) Stop location (center of mass) plotted in arena coordinates aligned to pellet location (x-axis reversed in trials where the food pellet was placed on the right). Large dots indicate location of food pellet (blue) and sham pellet (grey). Grey outlines indicate 1 cm increments from food pellet. (ii) Probability of observing stops as a function of distance from food or sham pellet (KS test p = 2.005 e-11) (iii) Within 6cm radius, average stop distance relative to food pellet and sham pellet (rank sum p = 0.00001), black lines indicate median. b. Stop features for intact animals without food. (i) Stop location (center of mass) plotted in arena coordinates aligned to pellet location (x-axis reversed in trials where the food pellet was placed on the right). Large dots indicate location of food pellet (blue) and sham pellet (grey). Grey outlines indicate 1 cm increments from food pellet. (ii) Probability of observing stops as a function of distance from food or sham pellet (KS test p = 0.007) (iii) Within 6cm radius, average stop distance relative to food pellet and sham pellet (rank sum p = 0.4650), black lines indicate median. c. Stop features for full antenna removed animals with food (i) Stop location, as in (a) (ii) Probability of observing stops as a function of distance from food or sham pellet (KS test p = 2.53 e-13) (iii) Within 6cm radius, average stop distance relative to food pellet and sham pellet (rank sum p =.2233), black lines indicate median. d. Stop features for first antenna removed animals with food (i) Stop location, as in (a) (ii) Probability of observing stops as a function of distance from food or sham pellet (KS test p = 6.16 e-13) (iii) Within 6cm radius, average stop distance relative to food pellet and sham pellet (rank sum p =.1579), black lines indicate median. e. Stop features for second antenna removed animals with food (i) Stop location, as in (a) (ii) Probability of observing stops as a function of distance from food or sham pellet (KS test p =.00037) (iii) Within 6cm radius, average stop distance relative to food pellet and sham pellet (rank sum p =.0943), black lines indicate median.

## Notes

### Competing Interest Statement

The authors have declared no competing interest.

