## Supplementary figures and images for "Ballistic food approaches in *Parhyale hawaiensis* require the antennae"

### Supplementary Video 1

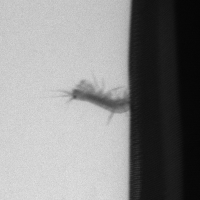
